# Manipulating speech perception with amplitude-modulated kilohertz magnetic perturbation (AM-kTMP)

**DOI:** 10.64898/2026.07.17.739132

**Authors:** Ram K. Pari, Christina Merrick, Philipp Reber, Cidnee Luu, Richard B Ivry, Daniel Sheltraw, Ludovica Labruna, Benedikt Zoefel

## Abstract

Speech perception has been hypothesized to exploit neural activity that entrains to the rhythmic properties of the utterance. Perceptual sensitivity varies with the phase of oscillatory activity induced by speech, and with an external oscillatory perturbation induced by transcranial alternating current stimulation (tACS) over the temporal lobe. Here we used a new form of non-invasive brain stimulation (NIBS), kilohertz transcranial magnetic perturbation (kTMP), in which a low frequency perturbation is created by the amplitude modulation (AM) of a continuous high-frequency carrier signal (3.5 kHz). As a magnetic induction method, kTMP can produce much higher electrical fields at the cortical surface than tACS and reduces cutaneous co-stimulation. Human participants (n=40) listened to sequences of rhythmic speech while receiving kTMP with an AM component of 3.125 Hz. The phasic modulation of perceptual sensitivity was greater during AM-kTMP than during sham stimulation, suggesting that AM-kTMP entrains neural activity underlying speech perception. Participants’ ratings of rhythmic sensations and other side effects did not differ between AM-kTMP and sham stimulation. Together with its reduced peripheral stimulation, our results highlight kTMP as a promising NIBS method for modulating frequency-specific brain dynamics in speech perception.

## Introduction

The phase of spontaneous brain oscillations can modulate human perception (1–3). Rhythmic stimulation plays an important role in this context, as it provides a temporal scaffold to shape neural activity (4). Stimulus-brain synchronisation is often called “neural entrainment” (5,6). Due to a remarkable resemblance between the frequency of brain oscillations and rhythms found in human speech (7,8), it has been proposed that the entrainment of brain oscillations is fundamental for speech understanding (9,10). Indeed, neural entrainment to speech rhythm becomes stronger when speech is intelligible (11–13). Several lines of evidence converge to suggest that neural entrainment to speech and other stimuli involves brain oscillations (13–17). However, as endogenous brain oscillations are difficult to identify during rhythmic stimulation (8,18,19), we do not make this assumption in the present study, and use the term “neural entrainment” to refer to stimulus-aligned neural activity (6).

Non-invasive brain stimulation (NIBS) methods have been used to manipulate neural entrainment and demonstrated its causal role in various cognitive functions. Varying the phase of transcranial alternating current stimulation (tACS) relative to rhythmic sequences of speech sounds alters speech processing, presumably by modulating the alignment of neural activity with the speech rhythm (20–27). Measures of speech perception continue to fluctuate briefly at the stimulation frequency after tACS offset (13,14). Such sustained rhythmic effects support an entrainment of endogenous brain oscillations (18). However, due to tolerability limits imposed by peripheral nerve stimulation, electric conduction methods, such as tACS, only produce weak electric fields of ~0.2-0.5 V/m on the cortical surface at conventional intensities (1-2 mA) (28). Moreover, given that much of the applied current is shunted by the skin, it has been suggested that cutaneous stimulation, rather than direct brain stimulation, contributes to the phasic modulation of speech processing (29).

Magnetic induction methods, such as transcranial magnetic stimulation (TMS), can produce stronger cortical stimulation than tACS and have been instrumental in establishing a causal role of the superior temporal gyrus (STG) in speech processing (27,30,31). While rhythmic TMS (rTMS) can entrain neural activity (32), the TMS pulses are discernible as auditory clicks and the somatosensory taps associated with each pulse can become uncomfortable when targeting STG due to activation of facial muscles (33–35). While peripheral sensations can be matched with appropriate active controls, the rhythmic nature of the sensations may interfere with that of the speech signal. Moreover, the pulsed stimulation waveform has power in a wide range of frequencies, limiting the potential for probing frequency-specific effects on neural activity.

Kilohertz transcranial magnetic perturbation (kTMP) has developed as an alternative NIBS method capable of producing subthreshold E-fields that are much higher than that possible with tACS (36). kTMP is a magnetic induction method in which an amplifier is used to drive a conventional TMS coil in a continuous manner. Due to power constraints, the carrier frequency must be in the kHz range to induce significant, subthreshold E-fields at the cortical surface. Previous work, including experiments with kTMP, has shown that neural activity can be modulated by kHz stimulation (37,38). Moreover, amplitude modulation (AM) of the kHz signal can be used to introduce a perturbation at physiologically relevant frequencies. kTMP reduces peripheral nerve stimulation and, with appropriate procedures to mask somatosensory and auditory co-stimulation, participants are unable to distinguish between active kTMP and sham stimulation (37).

Here we used AM-kTMP to modulate neural entrainment in the left STG, following a protocol that has been successful in modulating speech perception with tACS (25,26). Participants listened to and reported monosyllabic, degraded words that were presented at 3.125 Hz in rhythmic sequences of five words (**Fig. 1A, orange**). AM-kTMP was applied at 3.5 kHz with the carrier frequency amplitude-modulated at 3.125 Hz throughout the task (**Fig. 1A, blue**). In the active stimulation condition, the intensity of stimulation was set to produce a cortical E-field of 5.03 or 7.82 V/m; in the sham condition, this cortical E-field was reduced to 0 V/m. Critically, the speech sounds were presented at different phases of the 3.125-Hz AM component. We expected neural activity to entrain to the AM component of kTMP (39–43), resulting in perceptual sensitivity varying as a function of the phase between the words and AM-kTMP (Fig. 1B).

**Figure 1.**
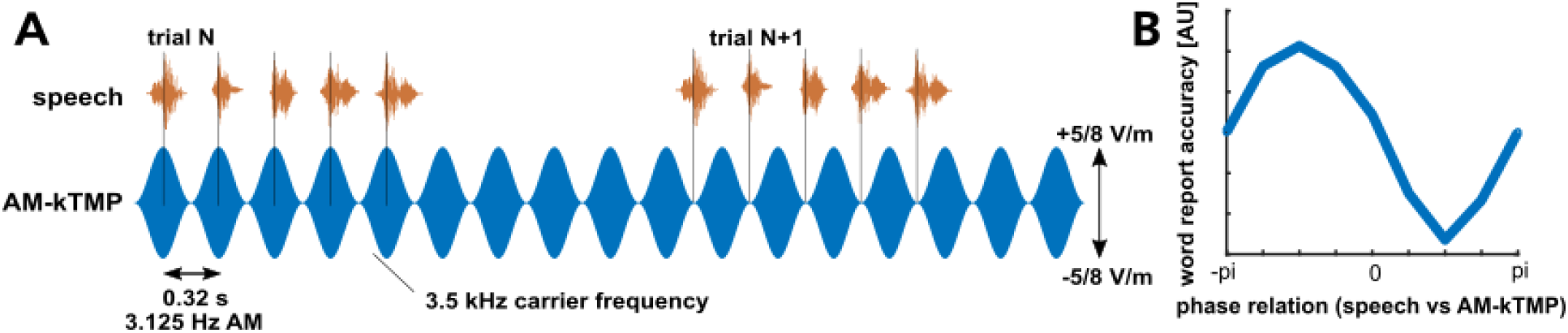
**A**. Experimental paradigm. Rhythmic sequences of monosyllabic spoken words (orange) were presented at different phases of the envelope of amplitude-modulated kTMP signal (blue). Participants were asked to report the words using a standard computer keyboard. **B**. We expected that accuracy in reporting words depends on the phase relation between speech and AM-kTMP. Note that the “best” phase (i.e., one associated with highest accuracy on speech perception) varies across individuals.

## Methods

### Participants

40 participants were recruited from the UC Berkeley community. Testing following the procedures outlined in a protocol approved by the IRB at UC Berkeley. All participants met inclusion criteria for research involving NIBS methods and provided informed written consent. The participants received $45 for their participation.

### Stimuli

The stimuli were developed in previous studies by our group (23,25,26). Monosyllabic words were initially spoken by a male speaker in time with a metronome beat (which was not presented to participants) at 1.6 Hz. The recordings were time-compressed offline to 3.125 Hz using the pitch-synchronous overlap and add (PSOLA) algorithm implemented in the Praat software (44). This compression ensured that stimulus rate corresponds to one where auditory sensitivity is relatively high (45) and might therefore produce relatively strong neural entrainment.

To create word sequences for each trial, the five words were combined to form sequences of rhythmic speech (**Fig. 1A, orange**). The first and the last words were always “pause”, included to serve as filler items. The middle three words were the target items. They were randomly selected from the stimulus set and linguistically unrelated to one another. The intelligibility of the words was reduced by embedding them in noise that spectrally matched the average spectral content of all words (4:1 ratio of root-mean squared sound amplitudes between speech and noise). The stimuli were presented to participants through insert earphones (ER-1, Etymotics).

### kTMP

The kTMP system (for details, see (37)) includes an amplifier that drives a coil to induce magnetic stimulation. In the present study, we used a Gen3 kTMP system (Magnetic Tides Inc., Berkeley, CA, USA) to drive a MagVenture Cool B-70 figure-of-eight coil (MagVenture A/S, Farum, Denmark). The carrier frequency was set to 3.5 kHz, and this signal was amplitude modulated at 3.125 Hz (**Fig. 1A, blue**). Stimulation was targeted at the left superior temporal gyrus (STG), based on standard MNI coordinates (62 −8 −2). To locate this position, we used the neuronavigation system Brainsight (Rogue Research, Montreal, Canada). The coil orientation was perpendicular to the target region (**Fig. 2A**), with the coil handle pointing upwards. This orientation minimized the distance between the coil and the scalp while avoiding contact with the ear. Coil position was monitored continuously with the Brainsight system, and the experimenter adjusted the position to offset subtle head movements by the participant.

**Figure 2.**
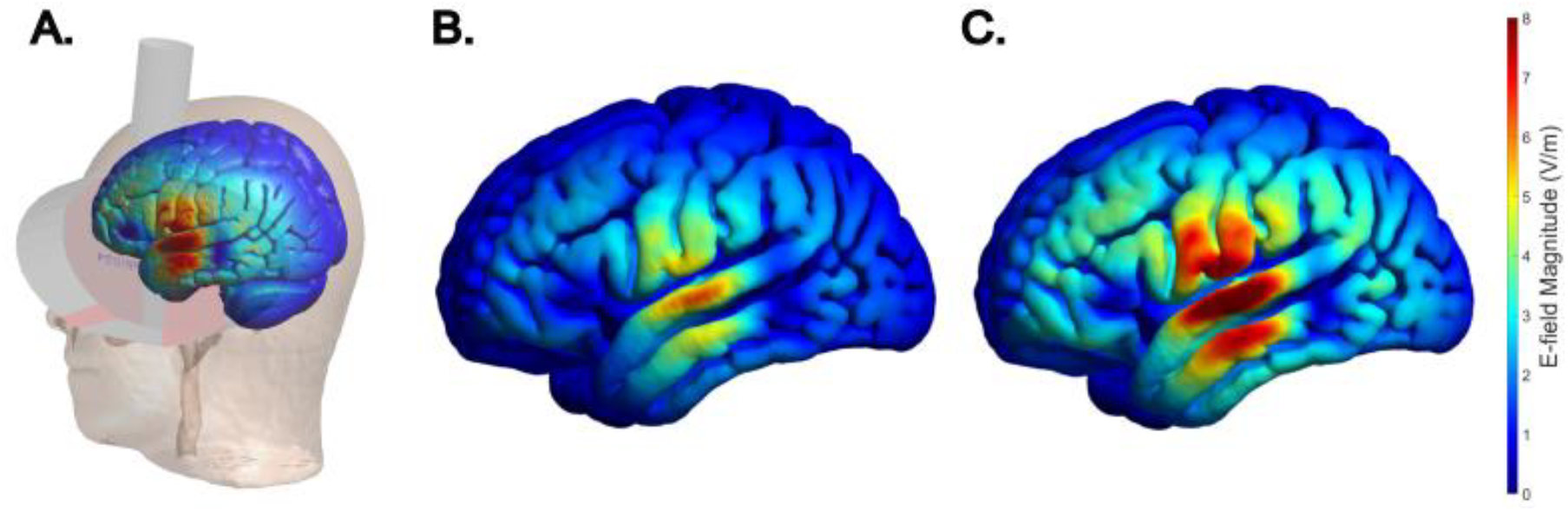
Electric field (E-field) during AM-kTMP, estimated using SIMNIBS with the coil positioned to target the left STG. **A**. Position of the TMS coil over the left STG. **B**,**C**. Simulated E-field distribution in which the current was set to produce maximal field strengths at left STG of ~5.03 V/m (B, first 20 participants) and ~7.82 V/m (C, second 20 participants), respectively.

We used the MNI152 template in the SIMNIBS 4.0 software (46,47) to simulate the actual E-field induced by AM-kTMP on the cortical surface, given a rate of change of current in the coil at 5.41 x 10^6^ A/s / 8.42 x 10^6^ A/s, and a coil distance of 10.5 mm from the scalp (6.5 mm default distance and additional 4 mm for a towel—see below). Based on these simulations, the maximum E-field at left STG was ~5.03 V/m for the first 20 participants (**Fig. 2B**). Following modifications, we were able to increase the maximum target field strength of the kTMP system and thus, the intensity was ~7.82 V/m for the second set of 20 participants (**Fig. 2C**). For sham stimulation, the AM-kTMP parameters remained the same as for real stimulation, but the intensity of the amplifier output was set to produce an E-field of 0 V/m.

We took two steps to mask peripheral sensations associated with kTMP. First, the system emits sound that contains a frequency component at the carrier frequency of stimulation and that is amplitude modulated during AM-kTMP. To mask this sound, we created an auditory mask to match its spectral profile and was played via loudspeakers located behind participants at approximately 20 dB louder than the kTMP-generated sound, corresponding to a 100x increase in physical intensity and a 4x increase in perceptual loudness. The mask was played during both AM-kTMP and Sham stimulation. Second, at the target scalp position used here, a slight vibration can be experienced from the coil during AM-kTMP. To mask this sensation, a towel was placed between the coil and the scalp. These procedures were successful in masking these peripheral sensations, as average sensations (including rhythmic ones) were consistently rated as weaker than “just noticeable” for both active AM-kTMP and sham stimulation, without reliable differences between the two (see Results).

### Experimental Paradigm

The experiment consisted of 8 blocks of ~4.2 minutes each. Half of the blocks involved real AM-kTMP stimulation and the other half involved sham stimulation. Block order was pseudo-randomised with the constraint that each condition occurred equally often across the experiment.

Each block consisted of 32 trials, with a trial defined as one five-word sequence (1.6 s long) and a subsequent response window. Within each trial, the five words were presented so that their perceptual centre (48), the part of the word that had been aligned with the metronome beat by the speaker, occurred at one of eight phases of the AM-kTMP waveform in steps of 45° (**Fig. 1B**). The phases were selected pseudo-randomly for each trial. Following the presentation of the sequence, the participant used a keyboard to type the middle three words. They were encouraged to guess or type in word fractions if unsure. The time window for participants’ responses was always 5.4 s, followed by an inter-trial interval that was variable in duration (0.4 s – 0.68 s) depending on the on the phase relation between speech and AM-kTMP on the trial just completed as well as the desired phase relation for the next trial. The phase relation between speech and AM-kTMP, along with corresponding inter-trial intervals, were also defined for sham stimulation.

After each experimental block (including sham), participants completed questionnaires about their perceived sensations from the stimulation on a scale of 0 to 10 (0 - No Sensation, 2 - Just Noticeable, 4 - Noticeable not aversive, 6 - Somewhat uncomfortable, 8 - Very Uncomfortable, 10 - Intolerable).

### Data analysis

Participants’ perceptual accuracy was quantified using Levenshtein distance (23), the minimum number of edits (deletions, insertions) necessary to change the response into the target word. To do so, responses and target words were converted into a phonetic representation using the “Phonemizer” package (49). This score was computed for each word and the score for the three words in a given trial were averaged. Given that the participant might shuffle the order when entering their responses, we calculated the Levenshtein distances for each combination of target and response words within a trial and selected the combination with the highest accuracy. Note that any bias introduced by this procedure will impact performance on AM-kTMP and sham trials.

We used a regression model to test how strongly the phase of the AM component of kTMP (sine- and cosine-transformed) predicts single-trial measures of performance (50). For individual participants, the model includes an intercept term and two circular predictors (one for sine, one for cosine predictor) that together reflect the magnitude of phasic modulation. This approach is equivalent to fitting a sine function to task accuracy as a function of oscillatory phase (**Fig. 1B**) and using the amplitude of that sine as an indicator of how strongly accuracy is modulated rhythmically. This amplitude corresponds to the degree AM-kTMP modulates perceptual accuracy in the word report task. Simulation analyses have shown that this method is optimal for detecting phase modulation (50).

We used the resulting regression coefficients to evaluate two hypotheses. First, we tested whether the phasic modulation in each stimulation condition (active AM-kTMP and sham stimulation) separately is reliably stronger than what would be expected under the null hypothesis. Here we tested the null hypothesis that all (sine- and cosine-based) coefficients except the intercept are zero in individual regression models, using an F-test (function “coefTest” in MATLAB). This comparison yielded one p-value per participant. This p-value is low if the regression model that includes AM-kTMP phase better predicts behavioural outcomes than a model that only includes an intercept term (but no phase). Individual p-values were combined to a group level p-value using Fisher’s method.

Second, we tested whether AM-kTMP produces a stronger phasic modulation of word report accuracy than sham stimulation. For this analysis, the two regression coefficients (sine- and cosine-based) were combined for each participant by calculating their root mean square, and a paired t-test (one-tailed, reflecting our directional hypothesis of larger coefficients for AM-kTMP) was used to compare the outcomes between AM-kTMP and sham stimulation. We used the Kolmogorov-Smirnov test to ensure that the assumption of normality in the distribution of paired differences could not be rejected. We also verified that observed effects were not driven by outliers by re-running the statistical test after exclusion of paired differences lying more than 2 standard deviations from the mean.

We also included an unplanned comparison of stimulation intensity, taking advantage of the modifications in the AM-kTMP system that allowed us to target a higher E-field for the second half of the participants. We contrasted the regression coefficients from the AM-kTMP condition for participants who received either ~5.03 V/m or ~7.82 V/m stimulation (two-sample t-test).

## Results

We first assessed whether AM-kTMP has generic effects on speech perception, independent of phase, comparing AM-kTMP and sham stimulation on word report accuracy (1 - Levenshtein distance; see Data Analysis). A paired t-test between the two conditions showed no significant difference (**Fig. 3A**, t(39) = −0.14, p = .553). In addition, word report accuracy did not differ between participants that received ~5.03 V/m stimulation and those that received ~7.82 V/m (t (38) = 1.03, p = .311).

**Figure 3.**
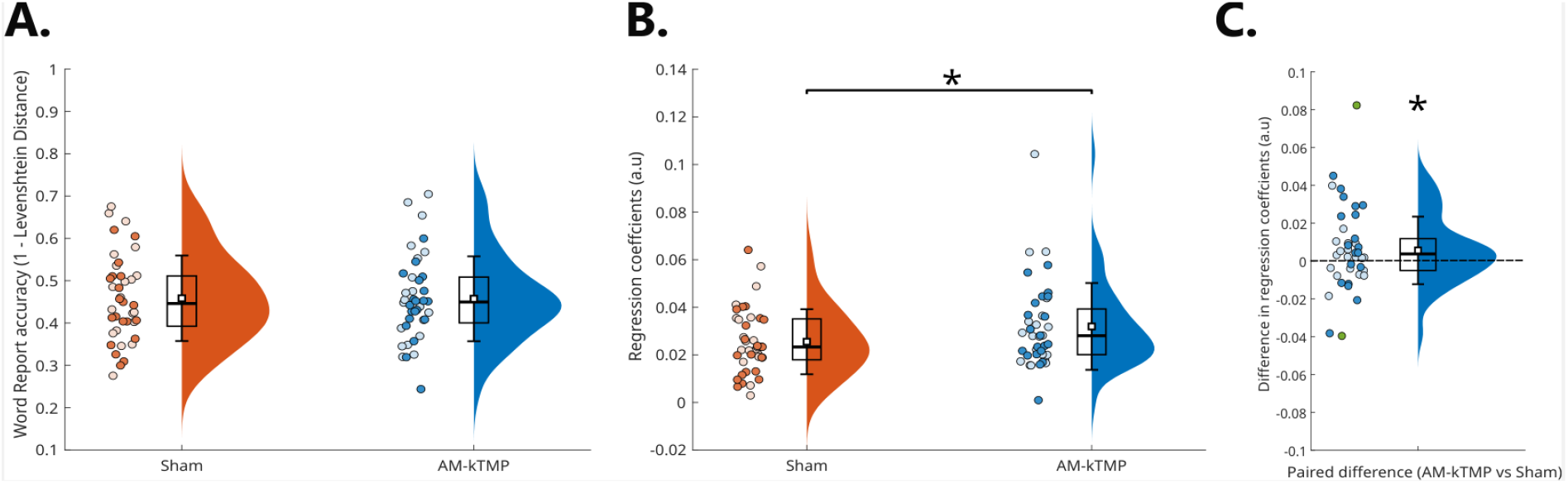
Effects of AM-kTMP on speech perception. **A**. Accuracy in reporting target words (1 - Levenshtein distance), averaged across the AM-kTMP/speech phases. Orange is sham stimulation, blue is AM-kTMP stimulation. **B**. Phase-dependent modulation of speech perception. Regression coefficients (root mean square of sine and cosine predictors) reflecting how accuracy is modulated by the phase relation between AM-kTMP/sham and speech. **C**. Paired differences between AM-kTMP and Sham conditions shown in B. Data points lying more than 2 standard deviations from the mean are indicated in green. In all panels, data from individual participants are shown as circles, where lighter and darker colours represent the stimulation intensity they received (5.03 or 7.82 V/m). Average (white square), standard deviation (whiskers), and percentiles (25, 50, 75th percentile – Box plot) are displayed along with visualizations of their distribution. Asterisk indicates a p-value < 0.05. The same statistical test underlies panels B and C, as the paired t-test in B corresponds to a test of the paired differences (AM-kTMP vs sham) against 0.

### AM-kTMP produces a stronger phasic modulation of speech perception than sham stimulation

Turning to our main hypothesis, we used a regression-based analysis to test whether speech perception was modulated as a function of the phase of kTMP’s AM component. We modulated the carrier frequency by the AM component in both the kTMP and sham conditions, although in effect, there was no modulation in the sham condition since the amplitude of the E-field was set to 0 V/m.

Our regression-based approach does not assume that the “best” phase (phase corresponding to most accurate speech perception) will be consistent across individuals; indeed, this heterogeneity was observed in related tACS work (50). **Fig. S1** shows data from example participants, selected to illustrate strong (left column), medium (middle column), and weak (right column) phasic modulations of word report accuracy. The amplitude of sine functions fitted to individual data (dotted lines in **Fig. S1**) is equivalent to the regression coefficients used for statistical analysis.

We first tested whether the phasic modulation in each stimulation condition (active AM-kTMP and sham stimulation) separately is reliably stronger than what would be expected under the null hypothesis. This was achieved by testing the null hypothesis that the (sine- and cosine-based) regression coefficients except the intercept are zero (F-test), yielding one p-value per participant. Individual p-values were combined to a single group-level p-value per condition with Fisher’s method. Phasic modulation was not significant in either the AM-kTMP condition (p = .175) and sham condition (p = .990). We note that, unlike the contrast of paired differences described next, this analysis does not exploit within-subject covariance.

We then used a paired t-test to contrast regression coefficients between AM-kTMP and sham stimulation. For this test, sine- and cosine-based coefficients were combined by calculating their root mean square, yielding values that are necessarily positive (**Fig. 3B**). We found higher combined regression coefficients during AM-kTMP stimulation than during sham stimulation (t (38) = 1.81, p = .039; Cohen’s d = 0.29), indicating a stronger phasic modulation during AM-kTMP than during sham. This statistical test is equivalent to comparing paired differences (AM-kTMP vs sham, **Fig. 3C**) to 0. While the paired differences met the assumption of normality (p = .27, Kolmogorov-Smirnov test), we verified that the exclusion of two outlier data points (green in **Fig. 3C**) did not change this outcome (t (36) = 1.95, p = .030). The significant difference between conditions suggests that AM-kTMP entrained neural activity, leading to weak, albeit reliable effects on speech perception.

Due to ongoing developments with the kTMP system, the second half of the participants received stronger stimulation than the first half, allowing us to perform an unplanned comparison of the effect of stimulation intensity. The phasic modulation during AM-kTMP did not differ between the two halves (t (38) = 0.49, p = .623).

### Sensations during AM-kTMP do not differ from sham stimulation

The participants provided reports of experienced sensations after each block. These reports are important to assess tolerance/comfort of AM-kTMP and evaluate if participants subjective experience differed between AM-kTMP and sham stimulation. We were especially interested in evaluating rhythmic sensations as the sound produced during AM-kTMP stimulation has an AM component which we sought to mask with a louder, non-AM sound. As shown in **Fig. 4**, all sensations including “rhythmic sensations” were rated at a level below “just noticeable”. There were no differences between AM-kTMP and sham stimulation in any of the categories (all p > .05; paired t-tests). These data are in accord with the results from a previous study indicating that kTMP can be employed in a manner that does not produce sensations that can reliably be distinguished from a sham condition (37). They also indicate that the masking sound was successful in blocking perception of the AM component emitted by the amplifier, and that the towel dampened the slight vibration produced by the coil at the current intensities used in this study.

**Figure 4.**
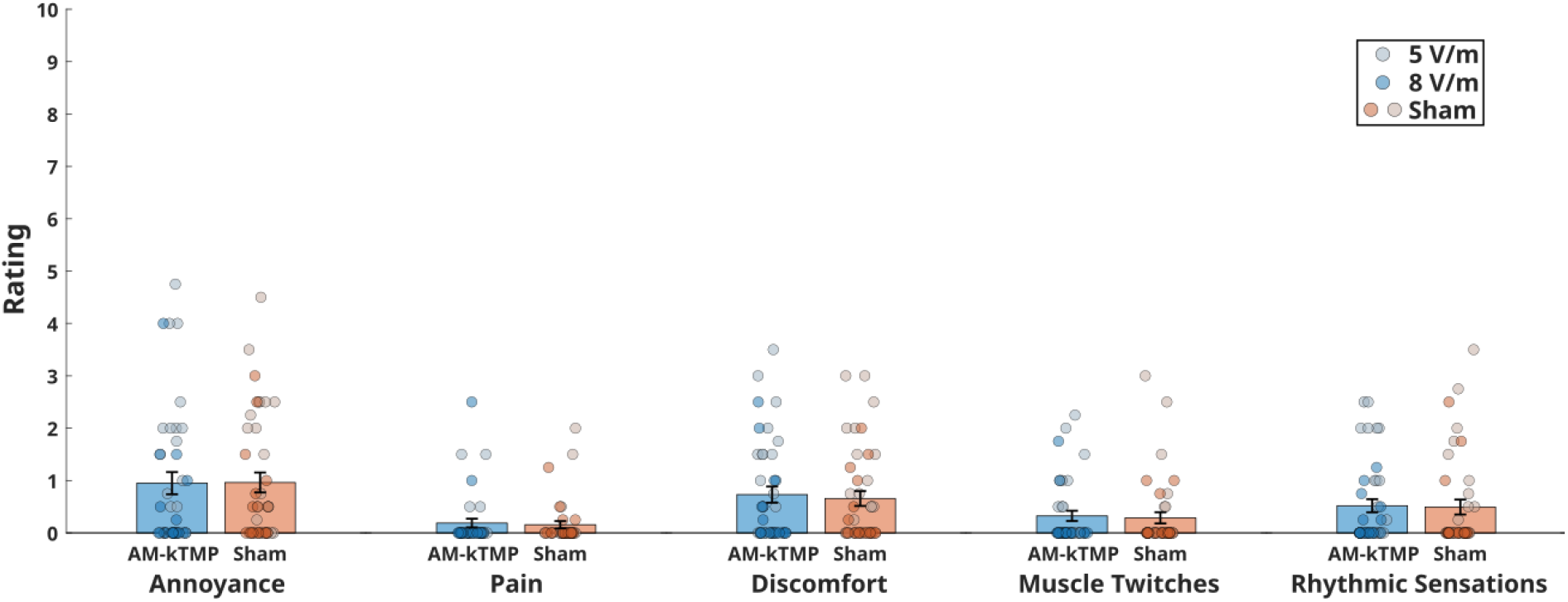
Sensation ratings on a 10-point scale (0 = No Sensation; 2 = Just Noticeable; 4 = Noticeable not aversive; 6 = Somewhat Uncomfortable; 8 = Very Uncomfortable; 10 = Intolerable) during AM-kTMP (blue) and sham stimulation (orange). The error bars indicate the Standard mean error (SEM). Data from individual participants are shown as circles, with lighter and darker circle colours based on the stimulation intensity they received.

## Discussion

We here used a novel transcranial magnetic stimulation technique, AM-kTMP, to modulate speech perception, putatively through a manipulation of neural entrainment. Our results indicate that AM-kTMP can modulate neural activity entrained to rhythmic properties of speech, and complement previous tACS work suggesting that this entrainment causally modulates speech processing (20–27,29).

A key drawback of NIBS is its sensory co-stimulation that can be auditory, visual (retinal), or somatosensory, depending on the protocol used. During tACS, a large portion of the current is shunted by the skin (51), which reduces direct brain stimulation and leads to simultaneous cutaneous stimulation that can affect neural or perceptual outcomes (52). Previous work suggested that cutaneous stimulation can indeed contribute to phasic tACS effects in speech perception (29). Conventional TMS produces considerable tactile sensations (53,54) as well as ‘click’ sounds that follow the stimulation rate during rTMS. As AM-kTMP applies magnetic (rather than electric) and continuous (rather than pulse-shaped) stimulation, and the sound or vibrations it produces can be masked, this new NIBS method strongly reduces cutaneous stimulation and associated sensations (**Fig. 4**). Sensations reported, including rhythmic ones, did not differ between AM-kTMP and sham stimulation, replicating a previous study (37). These observations highlight AM-kTMP as promising method to study the causal role of neural entrainment in different processes and regions. Nevertheless, the effect size observed here (Cohen’s d = 0.29 for the contrast between AM-kTMP and sham stimulation) resembles those typically obtained with tACS (22,23,26). Although this might seem surprising at first glance, given the significantly stronger electric field produced by kTMP, there are several points to consider.

First, AM-kTMP has only been developed recently. Consequently, we applied AM-kTMP for only ~16 minutes per participant, obtaining a lower number of trials compared to previous tACS work. It is likely that longer and stronger stimulation will become routine and improve statistical power in the future. Dose-response curves need to be established for AM-kTMP. The fact that phasic modulation did not differ between 5.03 V/m and 7.82 V/m indicates that the curve is relatively flat between these intensities. Due to its relatively young age, optimal parameters for the use of AM-kTMP to manipulate neural entrainment to speech also remain unknown – including the question whether the left STG is the ideal target for this purpose. The minimum stimulation intensity to achieve reliable effects in auditory cortical regions is yet to be established. For the motor and visual systems, such effects are often expressed relative to available correlates of neural excitability, such as the motor evoked potential or phosphenes (55,56), but auditory research lacks an equivalent marker. This makes it difficult to adjust stimulation intensities to individual response thresholds.

Second, improvements are possible to achieve consistent and efficacious stimulation effects in the target region. Electric field simulations should be combined with individual structural MRI scans (57) to extract target locations for individual participants. Cortical folding patterns (gyrification) represent a key factor of variability in non-invasive brain stimulation (58–60), possibly amplified by anatomical variation in the STG (61). The dorsoventral angle of current injection of our stimulation montage (illustrated in **Fig. S2**) might be sensitive to such variation, due to a high number of variability folds and their respective folding angles in the dorsoventral plane. Moreover, since the magnetically induced electric field is not fixed relative to the participant’s head, it is subject to variations due to head movement. Improvements in neuronavigation hardware and automated repositioning of the induction coil would be helpful in this regard.

Third, neural entrainment at physiologically relevant frequencies is assumed to be achieved through kTMP’s AM component. Previous tACS work suggest that entrainment effects are weaker if they are conveyed through an amplitude modulated carrier as compared to conventional (unmodulated low-frequency) tACS (41,62,63). However, unlike tACS, the kTMP electric field magnitude can, with hardware development, be safely increased by at least a factor of 5 or more before peripheral nerve or tissue heating becomes problematic. Such developments are expected to produce greater efficacy with respect to AM stimulation.

Several complementary or alternative stimulation protocols are plausible for the use of AM-kTMP to manipulate neural entrainment. These include an adaptation of the AM frequency to that of endogenous neural rhythms (64), the application of non-sinusoidal waveforms (65), or the use of bilateral stimulation protocols that have been shown to enhance tACS effects (26). Post-hoc sorting of stimulation phases can be inefficient, and a more targeted stimulation (e.g., exclusively of individual best or worst phases) can improve effect sizes. It has been reported that individual best tACS phases can be predicted from electroencephalographic (EEG) responses to rhythmic speech (14,23). If a similar correlation between EEG and AM-kTMP existed, it would simplify the use of AM-kTMP to manipulate entrainment. As the AM-kTMP signal does not contain power at the low frequencies presumed to entrain neural activity, a simultaneous use of EEG to is theoretically possible. Although additional work is required to remove artefacts caused by non-linear transfer characteristics (66,67), this would not only allow to measure entrained neural activity directly, but also a targeted (“closed-loop”) stimulation, relative to specific EEG phases (68,69).

Finally, temporal interference stimulation (TIS) relies on the simultaneous application of two or more stimulation waveforms to produce low-frequency electric fields in deeper brain regions (70). As continuous high-frequency waveforms are necessary for TIS that conventional TMS cannot deliver, TIS is currently applied through tACS (71). AM-kTMP can produce these waveforms, making it an interesting candidate method to apply magnetic TIS. Indeed, efforts to develop such protocols are ongoing (72).

## Conclusion

AM-kTMP is a novel form of NIBS that addresses several pressing issues in the field, including cutaneous co-stimulation and inflexible stimulation waveforms. We found that the phasic modulation of word report accuracy was stronger during AM-kTMP than during sham stimulation, suggesting that AM-kTMP entrained neural activity related to speech perception. Although effect sizes did not exceed those reported for more established methods, an optimisation and individualisation of AM-kTMP will likely increase effects further, and can make AM-kTMP a key method in the field of neural entrainment and its effect on cognition.

## Funding

This study has been supported by the Fyssen Foundation (BZ) and through the grant EUR CARe N°ANR-18-EURE-0003 in the framework of the Programme des Investissements d’Avenir (RKP, BZ). Support was provided by the National Institute of Health (Grants 1R44NS127667 (LL), R44NS139730 (LL), R35NS116883 (RBI)). Additional support was provided by the Weill Neurohub Next Great Ideas Fellowship and the Weill Neurohub Pillars Program (RBI, LL).

## Conflict of Interest

The authors declare the following competing interests: C.M., C.L., D.S., and L.L. are employees of Magnetic Tides Inc. P.R., C.M., C.L., L.L., D.S., and R.B.I. hold equity in Magnetic Tides. a non-publicly traded company created to develop new methods of non-invasive brain stimulation. The University of California, Berkeley holds patents related to the kTMP technology described in this study. The remaining authors declare no competing interests.

## Supplementary Materials

**Figure S1.**
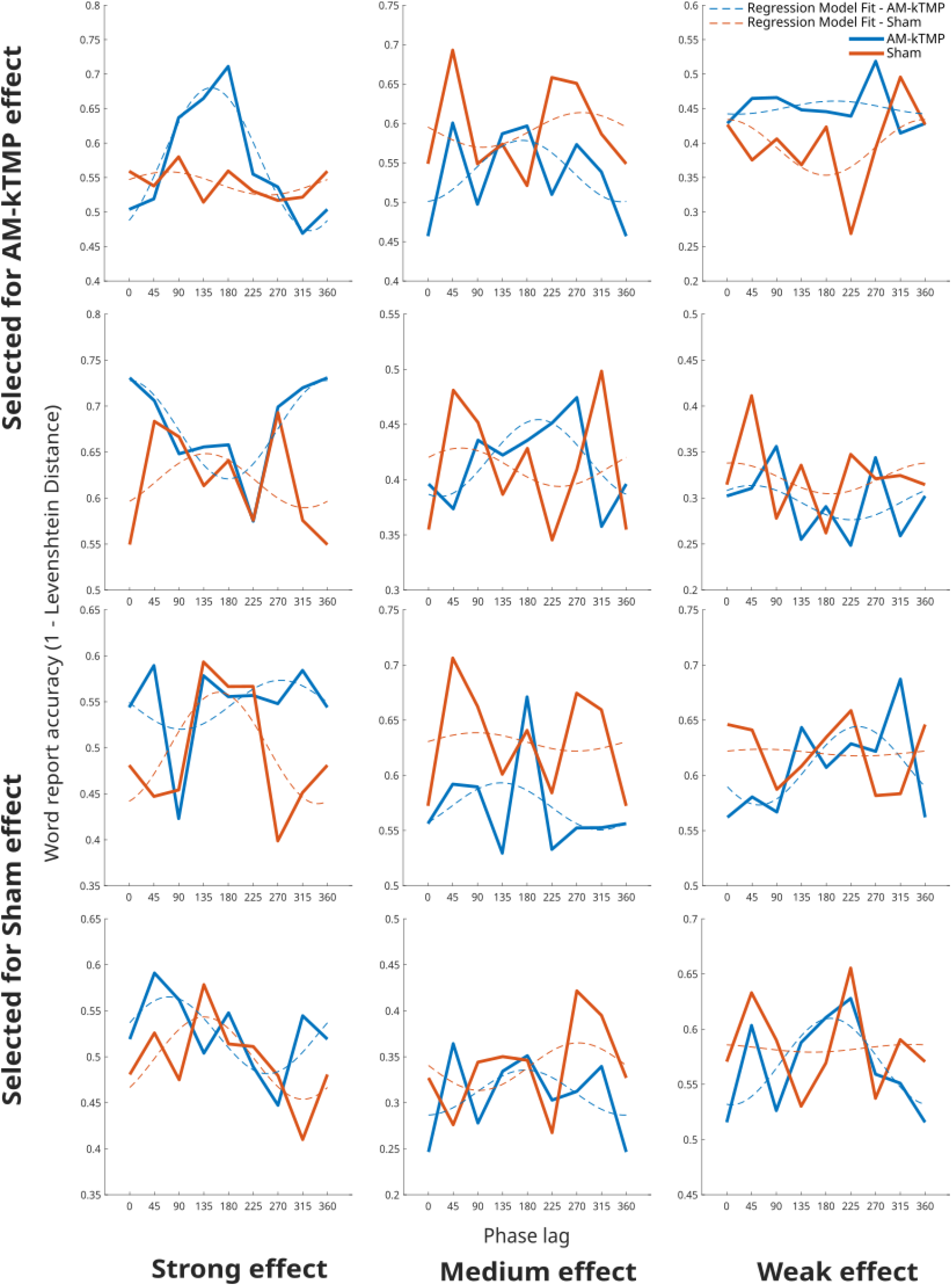
Word report accuracy as a function of phase for example participants. Participants have been selected to illustrate a relatively strong (left column), medium (middle column) and weak (right column) phasic modulation of word report accuracy in either the AM-kTMP (Blue) and Sham (Orange) condition. The dotted lines show the sinusoidal fits from the respective regression models.

**Figure S2.**
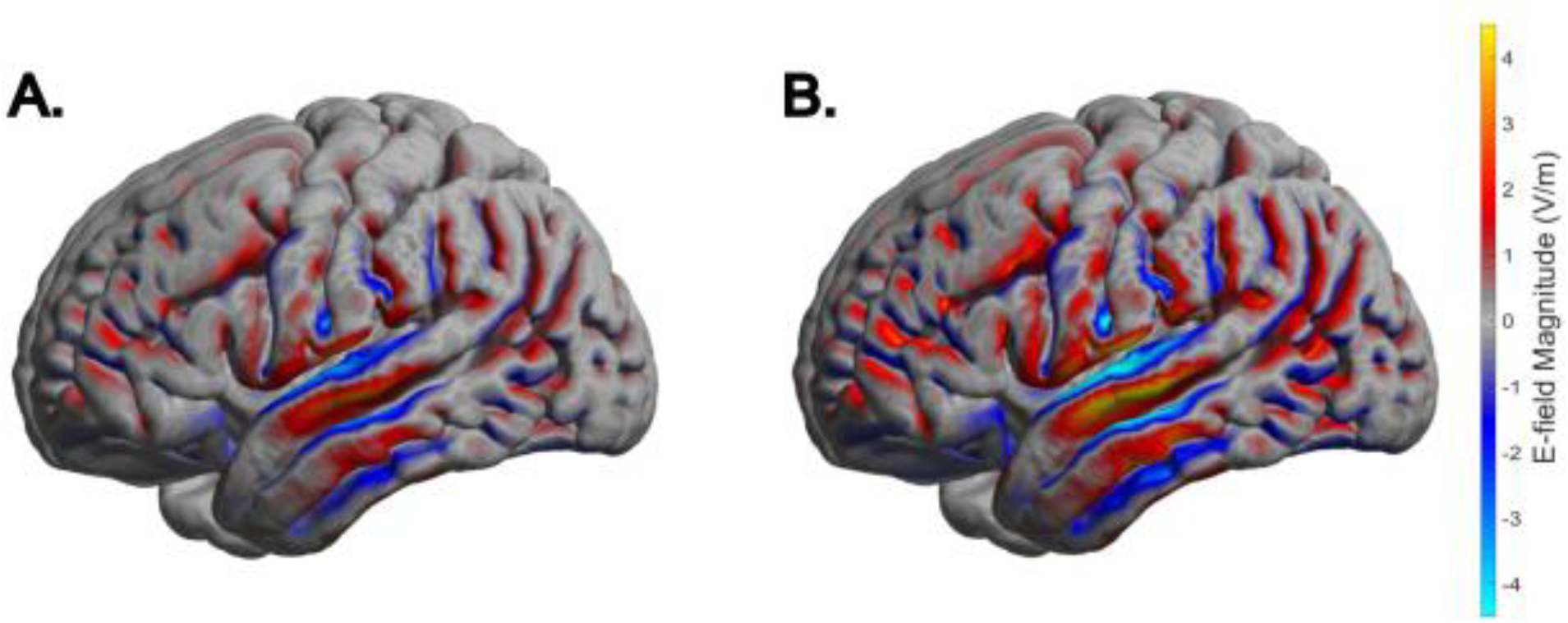
Normal component (the magnitude of the part that is perpendicular to the cortical surface) from the E-Field shown in Fig. 2B,C, estimated using SIMNIBS. The current was set to produce maximal field strengths (MagnE) at left STG of ~5.03 V/m (A, first 20 participants) and ~7.82 V/m (B, second 20 participants), respectively.

